# Next-generation sequencing identifies novel gene variants and pathways involved in specific language impairment

**DOI:** 10.1101/060301

**Authors:** Xiaowei Sylvia Chen, Rose H. Reader, Alexander Hoischen, Joris A. Veltman, Nuala H. Simpson, SLI Consortium, Clyde Francks, Dianne F. Newbury, Simon E. Fisher

**Affiliations:** Language and Genetics Department, Max Planck Institute for Psycholinguistics, Nijmegen, The Netherlands; Wellcome Trust Centre for Human Genetics, University of Oxford, Oxford, OX3 7BN; Department of Human Genetics, Radboud University Medical Center, Nijmegen, The Netherlands; Department of Clinical Genetics, University of Maastricht, Maastricht, The Netherlands; Donders Institute for Brain, Cognition and Behaviour, Nijmegen, The Netherlands; Department of Biological and Medical Sciences, Faculty of Health and Life Sciences, Oxford Brookes University, Oxford, UK

## Abstract

A significant proportion of children suffer from unexplained problems acquiring proficient linguistic skills despite adequate intelligence and opportunity. These developmental speech and language disorders are highly heritable and have a substantial impact on society. Molecular studies have begun to identify candidate loci, but much of the underlying genetic architecture remains undetermined. Here, we performed whole exome sequencing of 43 unrelated probands affected by severe forms of specific language impairment, followed by independent validations with Sanger sequencing, and analyses of segregation patterns in parents and siblings, to try to shed new light on the aetiology of the disorder. By first focusing on a pre-defined set of known candidates from the literature, we identified potentially pathogenic variants in genes already implicated in diverse language-related syndromes, including *ERC1, GRIN2A*, and *SRPX2*. Complementary analyses suggested novel putative candidate genes carrying validated variants which were predicted to have functional effects, such as *OXR1, SCN9A* and *KMT2D*. We also searched for potential “multiple-hit” cases; one proband carried a rare *AUTS2* variant in combination with a rare inherited haplotype affecting *STARD9*, while another carried a novel nonsynonymous variant in *SEMA6D* together with a rare stop-gain in *SYNPR*. When we broadened our scope to all rare and novel variants throughout the exomes, we identified several biological themes that were enriched for such variants, most notably microtubule transport and cytoskeletal regulation.

## INTRODUCTION

Developmental disorders of speech and language affect a large number of children and are related to educational, behavioural and psychological outcomes. Two primary language-related disorders that have been extensively investigated at the genetic level are specific language impairment (SLI) and developmental dyslexia. They impair spoken and written language skills respectively and are clinically defined as disorders affecting the given domain despite full access to education and no pre-existing neurological disabilities that might explain the impairment, such as an auditory or intellectual deficit (1). SLI and dyslexia are both highly heritable (2), and show high comorbidity, with complex genetic underpinnings involving multiple susceptibility loci (3). However, little is currently known regarding the crucial biological risk mechanisms.

A range of methods have been used to investigate the genetic architecture underlying speech and language disorders. Initial linkage studies of family-based samples identified SLI susceptibility loci on chromosomes 2p22, 13q21 (SLI3, OMIM%607134) (4), 16q23-24 (SLI1, OMIM%606711) (5), and 19q13 (SLI2, OMIM%606712) (5). Similarly, early studies of families affected by dyslexia uncovered regions of linkage on multiple chromosomes, including 15q21 (DYX1, OMIM#127700) (6), 6p22.3-p21.3 (DYX2, OMIM%600202) (7), 2p16-p15 (DYX3, OMIM%604254) (8), 3p12-q13 (DYX5, OMIM%606896) (9), 18p11.2 (DYX6, OMIM%606616) (10), 11p15.5 (DYX7) (11), 1p36-p34 (DYX8, OMIM%608995) (12) and Xq27.2-q28 (DYX9, OMIM%300509) (13). Subsequent investigations have identified associations and/or aetiological chromosomal rearrangements that implicate genes within several of these linkage regions (reviewed by (14)). Key genes include *CMIP* (C-maf-inducing protein, OMIM*610112) and *ATP2C2* (ATPase, Ca2+-transporting, type 2c, member 2, OMIM*613082) in SLI1 (15); *DYX1C1* (OMIM*608706) in DYX1 (16); *KIAA0319* (OMIM*609269) and *DCDC2* (Doublecortin domain-containing protein 2,OMIM*605755) in DYX2 (17-19); *C2orf3/MRPL19* (Mitochondrial ribosomal protein L19, OMIM*611832) in DYX3 (20); and *ROBO1* (Roundabout, Drosophila, homologue of, 1, OMIM*602430) in DYX5 (21). Additional risk loci and variations are beginning to be suggested by genome-wide association scans (GWAS, reviewed by (22)), but few have exceeded accepted thresholds for significance, and they have yet to be validated by independent replication studies.

Although the majority of speech and language impairments are modeled as complex genetic disorders, there is increasing evidence that common DNA variations are unlikely to provide a full account of their molecular basis (22). Thus, although linkage and association studies have identified strong evidence of a genetic influence, many rarer variants with aetiological relevance may be overlooked because they will not be captured by single nucleotide polymorphism (SNP) arrays, or do not reach stringent significance parameters. Recent findings indicate that the boundary between common traits and monogenic forms of disorder may be less defined than previously thought (23, 24). Accordingly, with the advent of next-generation sequencing, examples can be drawn from the literature of rare or private high-penetrance variants that contribute to certain forms of speech and language deficits (22). A study of an isolated Chilean population identified a coding variant within the *NFXL1* gene that was postulated to contribute to speech and language difficulties in this population (25). Mutations of the *FOXP2* transcription factor (Forkhead box, P2, OMIM*605317) are known to lead to developmental syndromes involving verbal dyspraxia, or childhood apraxia of speech, accompanied by problems with many aspects of language (26, 27). *FOXP1* (Forkhead box P1, OMIM*605515), a paralogue of *FOXP2*, has similarly been implicated in neurodevelopmental disorder (28, 29), along with some of its transcriptional targets, most notably, *CNTNAP2* (Contactin-associated protein-like 2, OMIM*604569) (30, 31). Rare variants of the *FOXP2* target *SRPX2* (Sushi-repeat-containing protein, X-linked, 2,OMIM*300642) have been identified in epileptic aphasias (32), as have mutations of *GRIN2A* (Glutamate receptor, ionotropic, N-methyl-D-aspartate, subunit 2A, OMIM*138253) (33, 34). Moreover, the closely related gene *GRIN2B* (Glutamate receptor, ionotropic, N-methyl-D-aspartate, subunit 2B, OMIM*138252) has also been implicated in language-relevant cognitive disorders (35–37). Overlaps between rare deletions and duplications that yield speech, language and/or reading disruptions have highlighted several additional candidate genes; including *ERC1* (ELKS/RAB6-interacting/CAST family member 1), *SETBP1* (SET-binding protein 1, OMIM*611060), *CNTNAP5* (Contactin-associated protein-like-5, OMIM*610519), *DOCK4* (Dedicator of cytokinesis 4, OMIM*607679), *SEMA6D* (Semaphorin 6D, OMIM*609295), and *AUTS2* (Autism susceptibility candidate 2) (38-44). Overall, this body of work points to the importance of rare and/or private variants in language-related phenotypes, suggesting that high-resolution molecular technologies like next-generation DNA sequencing hold considerable promise for unraveling a disorder such as SLI.

Thus, in this study, we performed exome sequencing of 43 probands affected by severe language impairment without a known cause. We employed complementary hypothesis-driven approaches to identify putative aetiological variants and associated biological processes. Our investigation detected cases with potential pathogenic mutations, and highlighted molecular pathways that may be important to speech and language development.

## RESULTS

### Exome sequencing in SLI

We performed whole exome sequencing of 43 unrelated probands affected by SLI (see *Subjects and Methods*). On average, 129.3 million mapped reads (median=133.3; min=67.1; max=173.3) were generated per sample. Across all 43 samples, an average of 85.5% of the target sequence was captured at a minimum depth of ten reads. The mean read depth of the exonic regions was 86.8, with 39.5% of reads reaching this level. Sequence metrics can be found in Supplementary Table S1. The coverage versus read depth of all samples is shown in Supplementary Figure S1.

In total, across all 43 probands, 353,686 raw variant calls were made, of which 62.2% fell outside known coding sequence. After removing variants with low quality (see *Subjects and Methods*), 270,104 remained. 35,550 (13.2%) of these were predicted to affect protein coding, including 34,571 nonsynonymous variants, 549 stop-gains/losses, and 430 splice-site variants. On average there were 8,594 (7,655-10,380) nonsynonymous variants, 91 (65-114) stop-gains/losses, and 72 (50-98) splice-site variants per individual (Supplementary Table S2).

The transition versus transversion ratio (Ti/Tv) for all SNVs within the exonic regions was 2.81, higher than the value observed for all variants (Supplementary Table S1), and in line with that expected (45). The total variants corresponded to 48,722 variants per individual (min=43,699; max=58,260) (Supplementary Table S1), the majority of which were common SNPs seen across all probands. As part of a prior published study (46), all 43 samples had previously been genotyped on Illumina HumanOmniExpress-12v1Beadchip (San Diego, CA, USA) arrays, which include ~750,000 common SNPs. 40,267 variants identified by our exome sequencing had been directly genotyped on the arrays and for these common SNPs,we observed a genotype concordance rate of 97%. The numbers of rare and novel variants identified per individual are shown in Supplementary Table S3.

In the first stage of analysis, we performed a tightly constrained search for aetiologically relevant variants, using several complementary methods. We began by identifying all variants occurring within a selection of known candidate genes that have previously been suggested as susceptibility factors in primary speech, language and/or reading disorders. Next, we characterized rare or novel variants of potential high risk from elsewhere in the exome by defining stop-gain variants, as well as searching for potential cases of compound heterozygotes for rare disruptive variants. Finally, we looked for likely “multiple-hit” events by searching for probands who carried more than one event of potential significance across different genes. For all variants in this stage of analysis we performed independent validation using Sanger sequencing, and assessed inheritance patterns in the available siblings and parents. Given the relatively small sample size of our study, these constraints provide a framework to maximize our chances of identifying contributory variants under an assumption that those variants will explain a large proportion of the trait variance. Throughout this paper, we refer to guidelines for inferring likely causality, as proposed by MacArthur and colleagues (47).

In the second stage of analysis, we broadened our scope to consider all rare and novel variants identified throughout the exome, and tested for biological pathways that showed enrichment in our dataset, using within-proband and group-based approaches. Moreover, we assessed how the pattern of findings was affected by the relative frequency of the variants being studied. Thus, this second stage went beyond the level of individual genes to provide a foundation for exploring potential mechanisms that could be involved in aetiology of SLI.

### Nonsynonymous variants in selected candidate genes

According to current guidelines for evaluating causality in whole exome/genome datasets, genes previously implicated in similar phenotypes should be evaluated before exploring potential new candidates (47). Therefore, prior to beginning any bioinformatic analyses of our exome data, we performed a literature search to identify a set of candidate genes that had been most reliably implicated in speech, language and reading disorders by earlier research. This literature survey yielded 19 candidate genes: *CMIP, ATP2C2, CNTNAP2* and *NFXL1*, which have previously been associated with common forms of SLI (15, 25, 30); *FOXP2*, which is involved in a monogenic form of speech and language disorder (26, 27), and its orthologue *FOXP1*, which has also been implicated in relevant neurodevelopmental disorders (28); *DYX1C1, KIAA0319, DCDC2, ROBO1*, which are candidate genes in developmental dyslexia (48); *SRPX2* and *GRIN2A*, which have been implicated in speech apraxia and epileptic aphasias (32, 34), as well as the closely related candidate *GRIN2B (36, 37)*; and, *ERC1, SETBP1, CNTNAP5, DOCK4, SEMA6D*, and *AUTS2*, each of which has been shown to have rare deletions or translocations that yield speech, language and/or reading disruptions (38-44).

We identified 37 coding or splice-site variants (36 SNVs, 1 insertion), that were successfully validated by Sanger sequencing, found in 14 of the 19 candidate genes (Table 1). A full list of these candidate-gene variants can be found in Supplementary Table S4. Seventy percent of validated calls represented common variants (population allele frequencies of >1% in 1000 genomes), 16.2% were rare variants (population frequencies <1% in 1000 genomes) and 13.5% represented novel changes (not present in 1000 genomes or EVS) (Table 1).

**Table 1.** Number of validated call id calls in candidate genes SLIC probands.

| **Gene** | Validated calls<br>with pop<br>freq >5%<sup>1</sup> | Validated calls<br>with pop freq<br>1-5%<sup>1</sup> | Validated calls<br>with pop freq<br><1%<sup>1</sup> | Novel<br>validated<br>calls<sup>2</sup> | Total<br>validated<br>calls |
| --- | --- | --- | --- | --- | --- |
| *ATP2C2* | 2 | 2 | 2 | 0 | 6 |
| *AUTS2* | 1 | 0 | 1 | 0 | 2 |
| *CMIP* | 0 | 0 | 0 | 0 | 0 |
| *CNTNAP2* | 0 | 0 | 0 | 1 | 1 |
| *CNTNAP5* | 0 | 1 | 1 | 0 | 2 |
| *DCDC2* | 2 | 1 | 0 | 0 | 3 |
| *DOCK4* | 0 | 0 | 0 | 0 | 0 |
| *DYX1C1* | 0 | 0 | 0 | 0 | 0 |
| *ERC1* | 1 | 1 | 0 | 1 | 3 |
| *FOXP1* | 0 | 0 | 0 | 0 | 0 |
| *FOXP2* | 0 | 0 | 0 | 0 | 0 |
| *GRIN2A* | 0 | 0 | 0 | 1 | 1 |
| *GRIN2B* | 0 | 0 | 0 | 1 | 1 |
| *KIAA0319* | 4 | 2 | 0 | 0 | 6 |
| *NFXL1* | 1 | 0 | 0 | 0 | 1 |
| *ROBO1* | 0 | 1 | 1 | 0 | 2 |
| *SEMA6D* | 2 | 0 | 0 | 1 | 3 |
| *SETBP1* | 5 | 0 | 0 | 0 | 5 |
| *SRPX2* | 0 | 0 | 1 | 0 | 1 |
| **All** | 18 (48.6%) | 8 (21.6%) | 6 (16.2%) | 5 (13.5%) | 37 |

1 – population frequency is taken from 1000 genomes (Apr2012\_ALL) samples
2 – novel variants were not described by 1000 genomes (Apr2012\_ALL) or by the exome variant server (EA5400\_ALL).
A full list of all 37 variants can be found in Supplementary Table S4.

In total, we observed 5 novel variants (in *ERC1, GRIN2A, GRIN2B, CNTNAP2* and *SEMA6D*) and 6 rare SNVs (in *ATP2C2, AUTS2, CNTNAP5, ROBO1* and *SRPX2*) in the predefined set of candidate genes (Table 2). All of these variants led to nonsynonymous changes. Those with an EVS European American allele frequency of <1% (n=9) were subsequently sequenced in available relatives to examine their segregation within the nuclear families (Figure 1, Supplementary Figure S2). Three such variants were considered the most likely to represent pathogenic changes based upon their inheritance, position in the protein and findings from previous literature. These include a *de novo* substitution (p.G688A) in a sporadic case in *GRIN2A* (with true *de novo* status validated via SNP data), a start-loss (disruption of the first methionine codon) in *ERC1* and a substitution (p.N327S) in *SRPX2* (Figure 1). We also observed a novel substitution in *SEMA6D* (p.H807D), and rare nonsynonymous changes in *AUTS2* (p.R117C) and *ROBO1* (p.V234A) that co-segregated with disorder in affected relatives of the respective probands (Supplementary Figure S2).

**Table 2.**
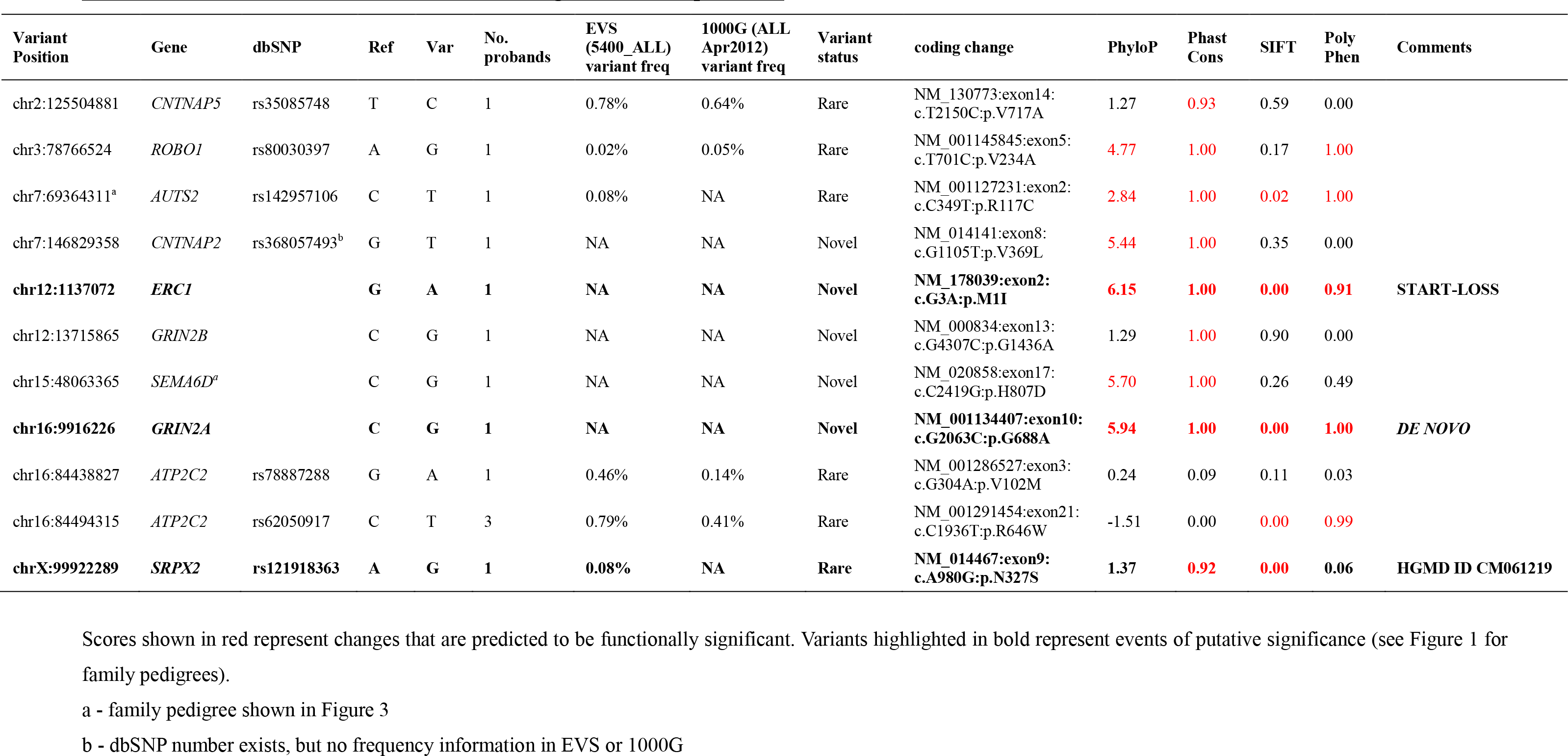
Novel and rare variants in candiate genes in SLIC probands

| Variant Position | Gene | dbSNP | Ref | Var | No. probands | EVS (5400_ALL) variant freq | 1000G (ALL Apr2012) variant freq | Variant status | coding change | PhyloP | Phast Cons | SIFT | Poly Phen | Comments |
| --- | --- | --- | --- | --- | --- | --- | --- | --- | --- | --- | --- | --- | --- | --- |
| chr2:125504881 | *CNTNAP5* | rs35085748 | T | C | 1 | 0.78% | 0.64% | Rare | NM_130773:exon14: c.T2150C:p.V717A | 1.27 | 0.93 | 0.59 | 0.00 |  |
| chr3:78766524 | *ROBO1* | rs80030397 | A | G | 1 | 0.02% | 0.05% | Rare | NM_001145845:exon5: c.T701C:p.V234A | 4.77 | 1.00 | 0.17 | 1.00 |  |
| chr7:69364311<sup>a</sup> | *AUTS2* | rs142957106 | C | T | 1 | 0.08% | NA | Rare | NM_001127231:exon2: c.C349T:p.R117C | 2.84 | 1.00 | 0.02 | 1.00 |  |
| chr7:146829358 | *CNTNAP2* | rs368057493<sup>b</sup> | G | T | 1 | NA | NA | Novel | NM_014141:exon8: c.G1105T:p.V369L | 5.44 | 1.00 | 0.35 | 0.00 |  |
| **chr12:1137072** | ***ERC1*** |  | **G** | **A** | **1** | **NA** | **NA** | **Novel** | **NM_178039:exon2: c.G3A:p.M1I** | **6.15** | **1.00** | **0.00** | **0.91** | **START-LOSS** |
| chr12:13715865 | *GRIN2B* |  | C | G | 1 | NA | NA | Novel | NM_000834:exon13: c.G4307C:p.G1436A | 1.29 | 1.00 | 0.90 | 0.00 |  |
| chr15:48063365 | *SEMA6D<sup>c</sup>* |  | C | G | 1 | NA | NA | Novel | NM_020858:exon17: c.C2419G:p.H807D | 5.70 | 1.00 | 0.26 | 0.49 |  |
| **chr16:9916226** | ***GRIN2A*** |  | **C** | **G** | **1** | **NA** | **NA** | **Novel** | **NM_001134407:exon10: c.G2063C:p.G688A** | **5.94** | **1.00** | **0.00** | **1.00** | **DE NOVO** |
| chr16:84438827 | *ATP2C2* | rs78887288 | G | A | 1 | 0.46% | 0.14% | Rare | NM_001286527:exon3: c.G304A:p.V102M | 0.24 | 0.09 | 0.11 | 0.03 |  |
| chr16:84494315 | *ATP2C2* | rs62050917 | C | T | 3 | 0.79% | 0.41% | Rare | NM_001291454:exon21: c.C1936T:p.R646W | -1.51 | 0.00 | 0.00 | 0.99 |  |
| **chrX:99922289** | ***SRPX2*** | **rs121918363** | **A** | **G** | **1** | **0.08%** | **NA** | **Rare** | **NM_014467:exon9: c.A980G:p.N327S** | **1.37** | **0.92** | **0.00** | **0.06** | **HGMD ID CM061219** |

a - family pedigree shown in Figure 3
b - dbSNP number exists, but no frequency information in EVS or 1000G

**FIGURE 1.**
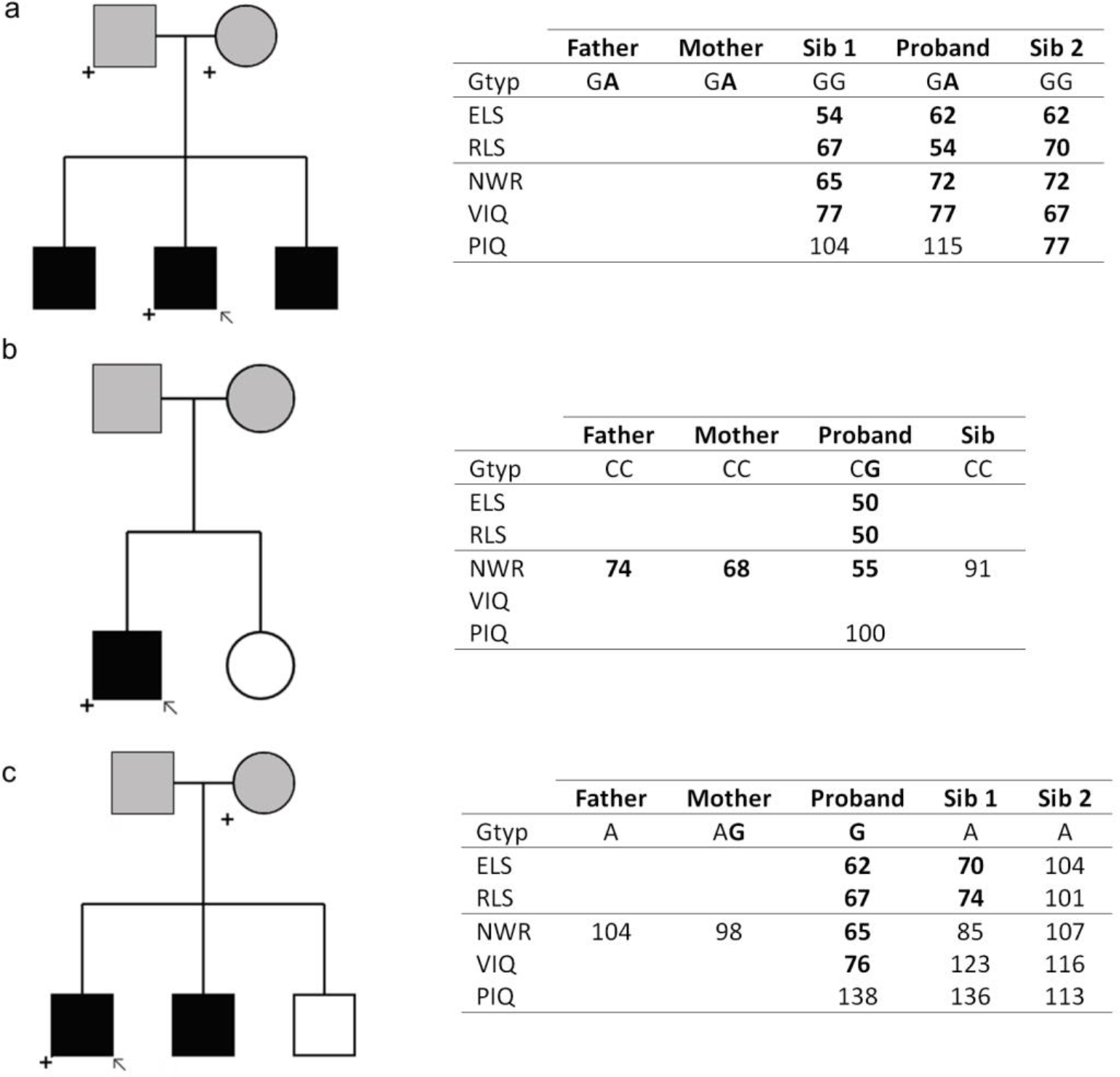
Variants of putative significance in candidate genes. ***1a — ERC1*. Chr12:1137072, NM_178039:exon2:c.G3A:p.M1I (start-loss)** Both parents report history of speech and language problems. All children have special educational needs. ***1b — GRIN2A*. Chr16:9916226, rs77705198, NM_001134407:exon10:c.G2063C:p.G688A (*de novo*)** Mother reports history of speech and language problems (although both parents have low NWR scores). Proband has special educational needs. ***1c — SRPX2*. ChrX:99922289, rs121918363. NM_014467:exon9:c.A980G:p.N327S** Parents do not report history of speech and language problems. All children have special educational needs. Proband is denoted by arrow. Individuals carrying variant allele are denoted by a plus symbol. Affected individuals are shaded black, unaffected are white, unknown are grey. Parents are always shaded as unknown as the language tests employed were for children only. Selfreported family history is given in text. Additional genotypic and phenotypic information is presented in inset table. Variant alleles are shown in bold. Affection status for all children was defined as CELF-R receptive (RLS) or expressive (ELS) language score >1.5SD below mean (see *Subjects and Methods* for details). We also present information regarding nonword repetition ability (NWR) and verbal and non-verbal IQ (VIQ and PIQ respectively). Although these additional scores were not used to ascertain affection status, they can provide useful information regarding specific deficits in individuals. NWR is thought to provide an index of phonological short term memory, while the IQ measures indicate a general level of verbal and non-verbal ability. All measures are standardized with a mean of 100 and a SD of 15. Scores >1.5SD below the mean are shown in bold.

### Variants of higher risk: rare stop-gains and potential compound heterozygotes

We next extended our investigation beyond known candidate genes, using two strategies to highlight coding variants of potential deleterious effect from elsewhere in the genome. In one approach, we identified and validated stop-gain variants in our dataset which are rare (< 1% in EVS and 1000 genomes) or novel. (We did not detect any validated rare/novel stop-loss or frame-shift variants in this dataset.) Stop-gain variants result in truncated proteins and have potential to yield more severe consequences than the majority of single amino-acid substitutions. In the other approach, we searched for genes that carried more than one rare, disruptive variant in the same proband, which may represent potential compound heterozygotes. (There were no instances where rare/novel disruptive variants occurred in the homozygous state in the cohort.) Within our sample, these approaches allowed us to focus upon variants that carry an increased chance of being deleterious. As recommended by MacArthur and colleagues (47), we targeted rare and novel variants, drawing upon large, ethnically matched control data and employing multiple bioinformatic prediction algorithms to evaluate potential pathogenicity. Moreover, again following accepted guidelines, we validated all variants of interest with an independent method (Sanger sequencing) and investigated co-segregation patterns within family units (47).

Following annotation and data filtering, we successfully validated 7 rare or novel stop-gain variants. These validated variants were found in the *OR6P1* (Olfactory receptor, family 6, subfamily P, member 1), *NUDT16L1* (Nudix (Nucleoside Diphosphate Linked Moiety X)-Type Motif 16-Like 1), *SYNPR* (Synaptoporin), *OXR1* (Oxidation resistance 1, OMIM*605609), *IDO2* (Indoleamine 2,3-dioxygenase 2, OMIM*612129), *MUC6* (Mucin 6, OMIM*158374) and *OR52B2* (Olfactory Receptor, Family 52, Subfamily B, Member 2) genes. Each was < 0.25% in reference samples and found to occur in a heterozygous state in a single proband in our dataset (Table 3). None occurred in known candidate genes for neurodevelopmental disorders. Note that olfactory receptor and mucin family genes are especially susceptible to false positive findings in next-generation sequencing, due to mapping artefacts (http://massgenomics.org/2013/06/ngs-false-positives.html). Thus, although these variants were validated by Sanger sequencing, they should be treated with caution. We again investigated the segregation of these variants within nuclear families (Supplementary Figure S3). Two variants showed evidence of co-segregation with disorder. One validated stop-gain, very near the start of the *OXR1* gene (NM_001198534:p.W5X, NM_001198535:p.W5X), was found in three children from a family, two affected by SLI necessitating special educational needs and a third with a diagnosis of dyslexia (Figure 2). The variant was not found in the mother, suggesting that it was most likely inherited from the father, who reports a history of speech and language difficulties but for whom we do not have any genetic information. In another pedigree, a validated stop-gain in *MUC6* (NM_005961:p.C703X) was passed from a father to four children, all of whom had expressive and receptive language difficulties (Figure 2).

**Table 3.** Stop gain variants identified in SILC probands.

| Variant Position | Gene | dbSNP | Ref | Var | No. probands | 1000G(ALL) variant freq | EVS (5400_ALL) variant freq | Variant status | Coding change | % of protein missing | PhyloP | Phast Cons | SIFT |
| --- | --- | --- | --- | --- | --- | --- | --- | --- | --- | --- | --- | --- | --- |
| chr1:158532597 | *OR6P1* | rs142215019 | G | T | 1 | 0.10% | 0.22% | rare | NM_001160325:exon1:<br>c.C798A:p.Y266X | 16.4% | -0.62 | 0.00 | 1.00 |
| chr3:63466576<sup>a</sup> | *SYNPR* | rs376661036 | C | A | 1 | NA | 0.01% | rare | NM_144642:exon2:<br>c.C93A:p.C31X | 84.7% | -0.72 | 0.60 | 1.00 |
| **chr8:107738486** | ***OXR1*** | **rs145739822** | **G** | **A** | **1** | **0.14%** | **NA** | **rare** | **NM_001198534:exon1:**<br>**c.G15A:p.W5X** | **97.7%** | **4.24** | **1.00** | **1.00** |
| chr8:39847306 | *IDO2* | rs199869245 | C | T | 1 | NA | 0.05% | rare | NM_194294:exon8:<br>c.C655T:p.R219X | 49.5% | 1.90 | 0.95 | 1.00 |
| **chr11:1027390** | ***MUC6*** | **rs200217410** | **G** | **T** | **1** | **NA** | **0.06%** | **rare** | **NM_005961:exon17:**<br>**c.C2109A:p.C703X** | **71.2%** | **0.40** | **1.00** | **1.00** |
| chr11:6190828<sup>a</sup> | *OR52B2* | rs190537696 | A | T | 1 | 0.14% | 0.10% | rare | NM_001004052:exon1:<br>c.T729A:p.C243X | 25.0% | 0.59 | 1.00 | 1.00 |
| chr16:4745030 | *NUDT16L1* | rs146701095 | C | T | 1 | 0.05% | 0.04% | rare | NM_001193452:exon3:<br>c.C556T:p.Q186X | 3.6% | 1.77 | 1.00 | 1.00 |

a – family pedigree shown in Figure 3

**FIGURE 2.**
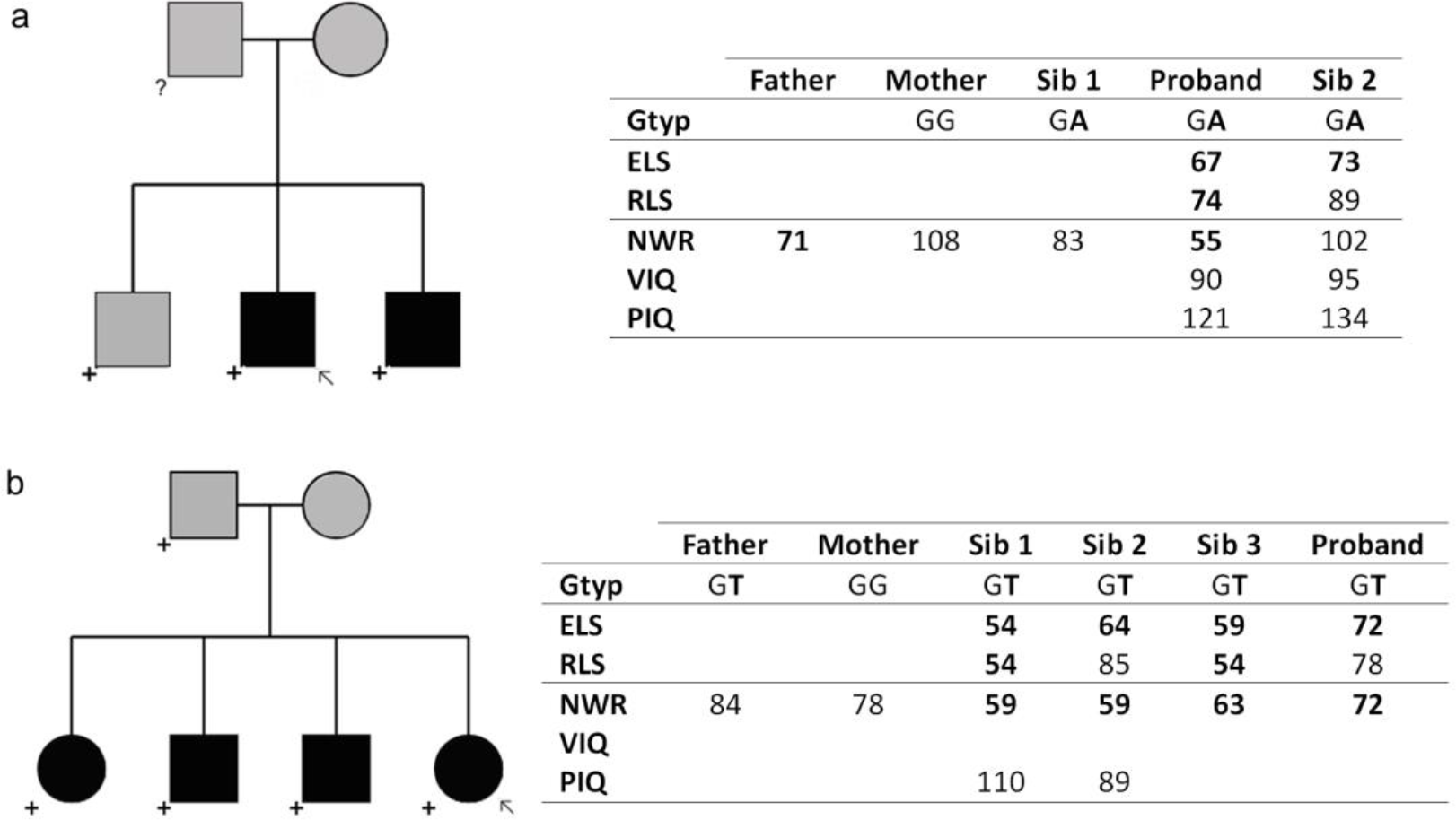
Co-segregating stop-gain variants. ***2a – OXR1*, Chr8:107738486, rs145739822, NM_001198534:exon1:c.G15A:p.W5X** Father reports history of speech and language problems. No DNA sample was available for father. Proband and sibling 2 have special educational needs. Sibling 1 does not have language or IQ scores available, but has been diagnosed with dyslexia. ***2b - MUC6*, chr11:1027390, rs200217410, NM_005961:exon17:c.C2109A:p.C703X** Mother reports history of speech and language problems. Proband has special educational needs. For key for symbols used in this figure, please refer to Figure 1.

In screening for potential cases of compound heterozygotes, we identified 11 genes which carried two or more rare or novel variants in the same proband (Table 4, Supplementary Figure S4). Upon family screening, four such cases were found to represent possible compound heterozygotes where two rare, potentially deleterious variants were inherited from opposite parents and co-segregated with disorder in the children (Supplementary Figure S4). The relevant variants occurred in the *FAT3* (Fat tumor suppressor, Drosophila, homologue of, 3, OMIM*612483), *KMT2D* (Histone-lysine N-methyltransferase 2D, OMIM*602113), *SCN9A* (Sodium channel, voltage-gated, type IX, alpha subunit, OMIM*603415) and *PALB2* (Partner and localizer of *BRCA2*, OMIM*610355) genes. Heterozygous mutations in the *SCN9A* gene have previously been associated with generalized epilepsy with febrile seizures (OMIM#613863) and Dravet syndrome (severe myoclonic epilepsy of infancy, OMIM#607208) when accompanied by mutations in the *SCN1A* (Sodium channel, neuronal type 1, alpha subunit, OMIM*182389) gene (49, 50). Loss-of-function mutations in *KMT2D* have been reported to cause Kabuki syndrome (OMIM#147920) (51-53), a severe syndromic form of intellectual disability associated with dysarthria and oromotor deficits, microcephaly and nystagmus (54). The *KMT2D* variants in our cohort were rare nonsynonmous changes, rather than confirmed loss-of-function mutations, and the individuals who carried them did not show features of Kabuki syndrome,

**Table 4.** Genes with more than one variant in the same SLIC proband

| Variant Position | Gene | dbSNP | Ref | Var | No. probands | 1000G(ALL) variant freq | EVS variant freq | Variant status | Coding change | PhyloP | Phast Cons | SIFT | Poly Phen | Notes |
| --- | --- | --- | --- | --- | --- | --- | --- | --- | --- | --- | --- | --- | --- | --- |
| **chr2:167089942** | **SCN9A** | **rs180922748** | **G** | **C** | **1** | **0.14%** | **0.20%** | **rare** | **NM_002977:exon21:c.C3799G:p.L1267V** | **2.07** | **1.00** | **0.13** | **0.99** |  |
| **chr2:167094638** | **SCN9A** | **rs141268327** | **T** | **C** | **1** | **0.41%** | **0.53%** | **rare** | **NM_002977:exon20:c.A3734G:p.N1245S** | **4.93** | **1.00** | **0.00** | **1.00** |  |
| chr2:32689842 | *BIRC6* | rs61754195 | C | T | 2 | 0.46% | 0.98% | rare | NM_016252:exon25:c.C5207T:p.P1736L | 3.09 | 0.99 | 0.01 | 0.60 |  |
| chr2:32740353 | *BIRC6* | rs61757638 | C | T | 2 | 0.18% | 0.25% | rare | NM_016252:exon55:c.C10865T:p.A3622V | 6.06 | 1.00 | 0.00 | 0.99 |  |
| chr3:58104626 | *FLNB* | rs139875974 | G | T | 1 | 0.05% | 0.07% | rare | NM_001164317:exon19:c.G2773T:p.G925C | 6.33 | 1.00 | 0.00 | 1.00 |  |
| chr3:58110119 | *FLNB* | rs111330368 | G | C | 1 | 0.23% | 0.41% | rare | NM_001164317:exon22:c.G3785C:p.G1262A | 6.18 | 1.00 | 0.00 | 1.00 |  |
| chr11:6190710 | *OR52B2<sup>a</sup>* | rs372373798 | C | T | 1 | NA | 0.01% | novel | NM_001004052:exon1:c.G847A:p.V283M | 2.69 | 1.00 | 0.08 | 1.00 |  |
| chr11:6190828 | *OR52B2<sup>ab</sup>* | rs190537696 | A | T | 1 | 0.14% | 0.10% | rare | NM_001004052:exon1:c.T729A:p.C243X | 0.59 | 1.00 | 1.00 | . | STOP |
| **chr11:92086828** | **FAT3** | **rs139595720** | **T** | **C** | **1** | **0.46%** | **0.66%** | **rare** | **NM_001008781:exon1:c.T1550C:p.L517S** | **3.32** | **0.82** | **0.72** | **1.00** |  |
| **chr11:92624235** | **FAT3** | **rs187159256** | **C** | **T** | **1** | **0.14%** | **0.17%** | **rare** | **NM_001008781:exon25:c.C13630T:p.L4544F** | **0.94** | **0.96** | **0.03** | **0.37** |  |
| **chr12:49418717** | **KMT2D<sup>a</sup>** | **rs201481646** | **C** | **T** | **1** | **NA** | **0.07%** | **rare** | **NM_003482:exon49:c.G15797A:p.R5266H** | **2.39** | **1.00** | **0.00** | **1.00** |  |
| **chr12:49432365** | **KMT2D<sup>a</sup>** | **rs199547661** | **G** | **A** | **1** | **0.09%** | **0.24%** | **rare** | **NM_003482:exon34:c.C8774T:p.A2925V** | **0.77** | **0.38** | **0.00** | **0.00** |  |
| chr13:109613971 | *MYO16* | rs374252281 | G | A | 1 | NA | 0.01% | rare | NM_001198950:exon18:c.G2122A:p.A708T | 4.62 | 1.00 | 0.01 | 1.00 |  |
| chr13:109617108 | *MYO16* |  | G | A | 1 | NA | NA | novel | NM_001198950:exon20:splice acceptor lost | 4.87 | 1.00 | . | . | SPLICE |
| chr14:58924684 | *KIAA0586<sup>a</sup>* | rs61742715 | T | A | 1 | 0.23% | 0.39% | rare | NM_001244189:exon13:c.T1729A:p.L577I | 0.28 | 0.81 | 0.43 | 1.00 |  |
| chr14:59014632 | *KIAA0586<sup>a</sup>* | rs61745066 | G | A | 1 | 0.18% | 0.24% | rare | NM_001244189:exon34:c.G4873A:p.G1625R | -1.15 | 0.41 | 0.00 | 0.00 |  |
| chr15:42977116 | *STARD9<sup>a</sup>* | rs79165890 | T | C | 1 | 0.05% | 0.22% | rare | NM_020759:exon23:c.T3340C:p.C1114R | 1.98 | 0.51 | 0.00 | 0.05 |  |
| chr15:42977810 | *STARD9<sup>a</sup>* | rs140924205 | T | G | 1 | 0.32% | 0.40% | rare | NM_020759:exon23:c.T4034G:p.I1345S | 0.269 | 0.001 | 0.00 | 0.27 |  |
| chr15:42978141 | *STARD9<sup>a</sup>* | rs376229251 | A | C | 1 | NA | 0.09% | rare | NM_020759:exon23:c.A4365C:p.E1455D | 0.553 | 0.067 | 0.00 | 0.99 |  |
| chr15:42981101 | *STARD9<sup>a</sup>* | rs202017657 | C | G | 2 | NA | 0.15% | rare | NM_020759:exon23:c.C7325G:p.P2442R | 0.28 | 0.01 | 0.00 | 0.99 |  |
| chr15:42982237 | *STARD9<sup>a</sup>* | rs201340789 | G | C | 2 | NA | 0.19% | rare | NM_020759:exon23:c.G8461C:p.V2821L | 0.28 | 0.01 | 0.00 | 0.99 |  |
| **chr16:23635348** | **PALB2** | **rs45478192** | **A** | **C** | **1** | **0.09%** | **0.17%** | **rare** | **NM_024675:exon8:c.T2816G:p.L939W** | **2.83** | **1.00** | **0.00** | **1.00** |  |
| **chr16:23641275** | **PALB2** | **rs45543843** | **T** | **A** | **1** | **NA** | **0.01%** | **rare** | **NM_024675:exon5:c.A2200T:p.T734S** | **2.90** | **0.97** | **0.11** | **1.00** |  |
| chr17:34861135 | *MYO19* | rs200572125 | C | T | 1 | NA | 0.03% | rare | NM_001163735:splice donor lost, exon20 | 4.98 | 1.00 | . | . | SPLICE |
| chr17:34871802 | *MYO19* | rs187710120 | T | C | 1 | 0.05% | 0.19% | rare | NM_001163735:exon8:c.A446G:p.Y149C | 4.52 | 1.00 | 0.00 | 1.00 |  |

Scores shown in red represent changes that are predicted to be functionally significant. Variants highlighted in bold represent potential compound heterozygotes.
a – family pedigree shown in Figure 3
b – stop-gain, also represented in Table 3

### Probands with multiple variants of putative interest

Four of the 43 probands investigated carried more than one rare variant across our prioritized high-risk categories, potentially representing “multiple-hit” events. The proband carrying a rare coding variant in *AUTS2* also had a stop-gain in *OR52B2*,and multiple rare variants in each of the *OR52B2, KIAA0586* (OMIM*610178) and *STARD9* (Start domain-containing protein 9, OMIM*614642) genes, all of which were successfully confirmed with Sanger sequencing. The majority of these variants were inherited from a mother who did not report a history of speech and language problems. Both siblings in this family were affected and both carried the rare variants in *AUTS2* and *STARD9* (Figure 3). Interestingly, in another family, a proband also carried multiple rare validated variants in the *STARD9* gene together with the rare missense variants in *KMT2D* mentioned above (Figure 3). In both families, the *STARD9* variants were not compound heterozygotes but instead appeared to represent inherited overlapping rare haplotypes that harboured multiple coding variants. One further proband carried a novel nonsynonymous variant in the *SEMA6D* (Semaphorin 6D, OMIM*609295) gene together with a rare stop-gain in the *SYNPR* gene (Figure 3). The proband is the only family member to inherit both variants and is the only family member with a history of speech and language impairment. Finally, one other family carried a novel variant in *GRIN2B* (Supplementary Figure S2) and two rare coding variants in *MYO19*. However, there was no obvious pattern of co-segregation across these variants.

**FIGURE 3.**
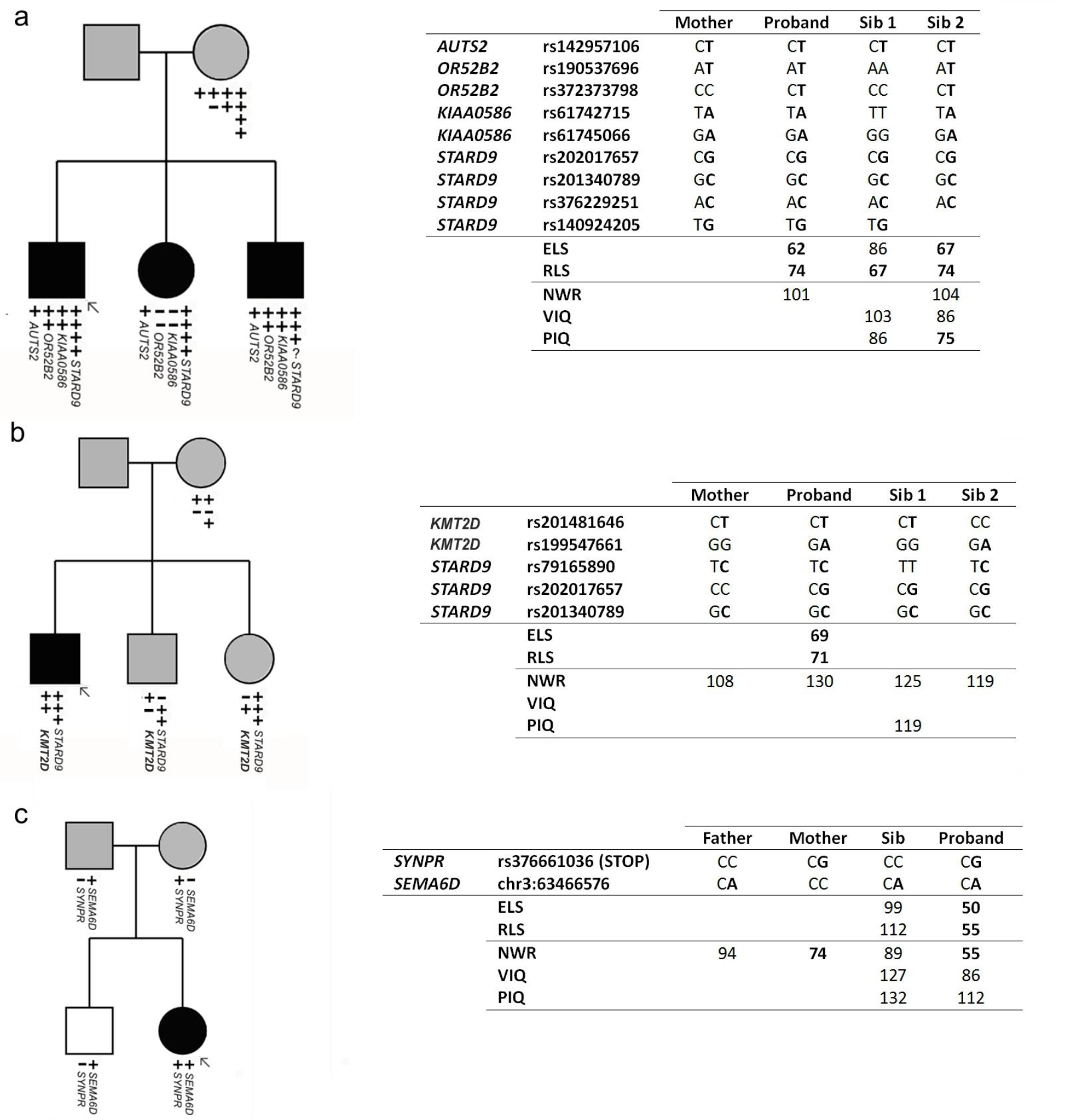
Probands with multiple hits of putative interest. **3a – Rare *AUTS2* variant, stop and rare variant in *OR52B2*, rare variants in *KIAA0586* and *STARD9*** Parents do not report history of speech and language problems. No sample available for father. All children have special educational needs. **3b – Multiple rare variants in *KMT2D* and *STARD9*** No family history available but maternal NWR score in normal range. No sample available for father. **3c – *SYNPR* rare stop variant and *SEMA6D* novel nonsynonymous variant** Parents do not report history of speech and language problems (although mother has low NWR score). Proband has special educational needs. For key for symbols used in this figure, please refer to Figure 1.

### Biological function enrichment analysis of genes with rare and novel SNVs

Prior studies suggest that, with a few prominent exceptions (27), most cases of speech and language impairments follow a complex disorder model where risk is determined by combinations of deleterious variants (55, 56). This is further supported by the observation of multiple rare events of potential significance in a subset of our families, described above. We therefore extended our studies to perform an exploratory exome-wide investigation that considered protein interaction pathways and networks. Although our sample is relatively small, these investigations are an important first step towards an unbiased assessment of the role of rare variants in SLI and will direct further studies in larger sample sets.

Within each proband, we generated a gene set corresponding to transcripts carrying rare (≤ 1% population frequency) or novel nonsynonymous SNVs, allowing the investigation of protein-interaction pathways within individuals. Pathways that were significantly shared by more than half of the probands included cell adhesion, regulation of the actin cytoskeleton, calcium signaling and integrin cell-surface interactions (FDR<0.01, Supplementary Table S5).

We went on to pool gene sets across all probands enabling the identification of gene ontology (GO) classes that were over-represented at the group level (with respect to nonsynonymous SNVs that are ≤ 1% or novel). The most significantly enriched GO term was GO:0001539: “ciliary of bacterial-type flagellar motility” (P=8.33x10^-5^), which is a small functional group consisting of 27 genes (Table 5). Twelve Dynein genes contributed to the 5-fold enrichment in this class. Other significantly over-represented terms included microtubule-based movement, cell adhesion, and actin cytoskeletal organization (FDR<0.01, Table 5).

**Table 5.** Enriched GO terms with variants less than 1% frequency across probands

| **Term** | **ExpCount** | **Count** | **P-value** | **FDR** |
| --- | --- | --- | --- | --- |
| **ciliary or bacterial-type flagellar motility** | 3.633861 | 12 | 8.33E-05 | 0.012 |
| **microtubule-based movement** | 20.45729 | 37 | 0.000197 | 0.009 |
| cell adhesion | 125.7047 | 160 | 0.000543 | 0.005 |
| homophilic cell adhesion | 18.70765 | 33 | 0.000683 | 0.008 |
| actin cytoskeleton organization | 55.04626 | 78 | 0.000777 | 0.004 |
| **extracellular matrix organization** | 46.56726 | 65 | 0.002975 | 0.011 |
| protein depolymerization | 8.748184 | 17 | 0.004587 | 0.001 |
| cellular component assembly involved in morphogenesis | 20.99564 | 33 | 0.005049 | 0.003 |
| dendrite development | 17.22719 | 28 | 0.005803 | 0.002 |
| double-strand break repair via homologous recombination | 6.998547 | 14 | 0.007348 | 0.004 |
| **neuromuscular junction development** | 4.979735 | 11 | 0.007624 | 0.005 |
| actin polymerization or depolymerization | 15.47756 | 25 | 0.00954 | 0.006 |
| cell projection organization | 126.5122 | 151 | 0.009759 | 0 |

In the pooled data set, we investigated the effects of expected variant frequency on pathway representation. We split our gene list into genes which carried novel variants and genes which carried variants that had been reported in the 1000 genomes and EVS with a variant frequency of <1%. To further test if genes carrying variants of a higher frequency (1%-5%) represented different pathway sets, we also included an additional set of genes with variants of expected frequency between 1% and 5%. Four related themes were found to be significant across variant frequency groups – microtubule-based movement, neuromuscular junction development, cilia and sequestration of calcium ions (Table 6). In general however, significant GO terms were found to cluster differently between frequency classes (Figure 4). Genes carrying potentially disruptive variants of higher frequency (1% to 5%) were predominantly localized within the classes “Cellular response to interleukin-4” and “Microtubule-based movement” while the GO enrichments for “Cell proliferation in forebrain” and “Extracellular matrix disassembly” relate mainly to the rarer variants (less than 1% and novel) (Figure 4).

**Table 6.** Enriched GO terms across probands split by various frequency

| Novel |  |  |  |  |
| --- | --- | --- | --- | --- |
| Term | ExpCount | Count | Pvalue | FDR |
| cellular response to interleukin-4 | 5.42 | 13 | 2.35E-04 | 0.006 |
| maintenance of location in cell | 24.07 | 40 | 2.41E-04 | 0.004 |
| keratinization | 3.85 | 10 | 9.66E-04 | 0.004 |
| **microtubule** anchoring | 7.94 | 15 | 0.0052 | 0.012 |
| **neuromuscular junction** development | 8.96 | 16 | 0.0078 | 0.003 |
| release of sequestered calcium ion into cytosol by sarcoplasmic reticulum | 4.09 | 9 | 0.0090 | 0.002 |
| Frequency between 0 and 1% |  |  |  |  |
| Term | ExpCount | Count | Pvalue | FDR |
| extracellular matrix disassembly | 23.67 | 41 | 8.57E-05 | 0.004 |
| cell proliferation in forebrain | 3.79 | 11 | 1.67E-04 | 0.004 |
| mitotic sister chromatid segregation | 11.36 | 23 | 2.06E-04 | 0.006 |
| **cilium** assembly | 20.12 | 34 | 2.16E-04 | 0.01 |
| **microtubule** anchoring | 7.34 | 16 | 6.78E-04 | 0.005 |
| cerebral cortex cell migration | 4.26 | 11 | 7.53E-04 | 0.003 |
| **neuromuscular junction** development | 8.29 | 16 | 0.0034 | 0.008 |
| **ciliary** or bacterial-type flagellar motility | 6.15 | 13 | 0.0031 | 0.004 |
| Frequency between 1% and 5% |  |  |  |  |
| Term | ExpCount | Count | Pvalue | FDR |
| **microtubule**-based movement | 46.38 | 76 | 1.81E-07 | 0.004 |
| homophilic cell adhesion | 45.69 | 74 | 3.26E-07 | 0.006 |
| **ciliary** or bacterial-type flagellar motility | 9.00 | 19 | 7.06E-05 | 0.005 |
| release of sequestered calcium ion into cytosol | 17.65 | 31 | 1.16E-04 | 0.006 |
| regulation of sequestering of calcium ion | 17.65 | 31 | 1.16E-04 | 0.006 |
| chemokine-mediated signaling pathway | 9.35 | 18 | 6.74E-04 | 0.004 |
| regulation of release of sequestered calcium ion into cytosol by sarcoplasmic reticulum | 4.73 | 10 | 0.0055 | 0.005 |
| sarcoplasmic reticulum calcium ion transport | 6.23 | 12 | 0.0053 | 0.003 |

Pathways with $P < 0.01$ and size $> 10$ genes are shown in the table. Highlighted key words indicated functions consistently found in all GO enrichment analysis.

**FIGURE 4.**
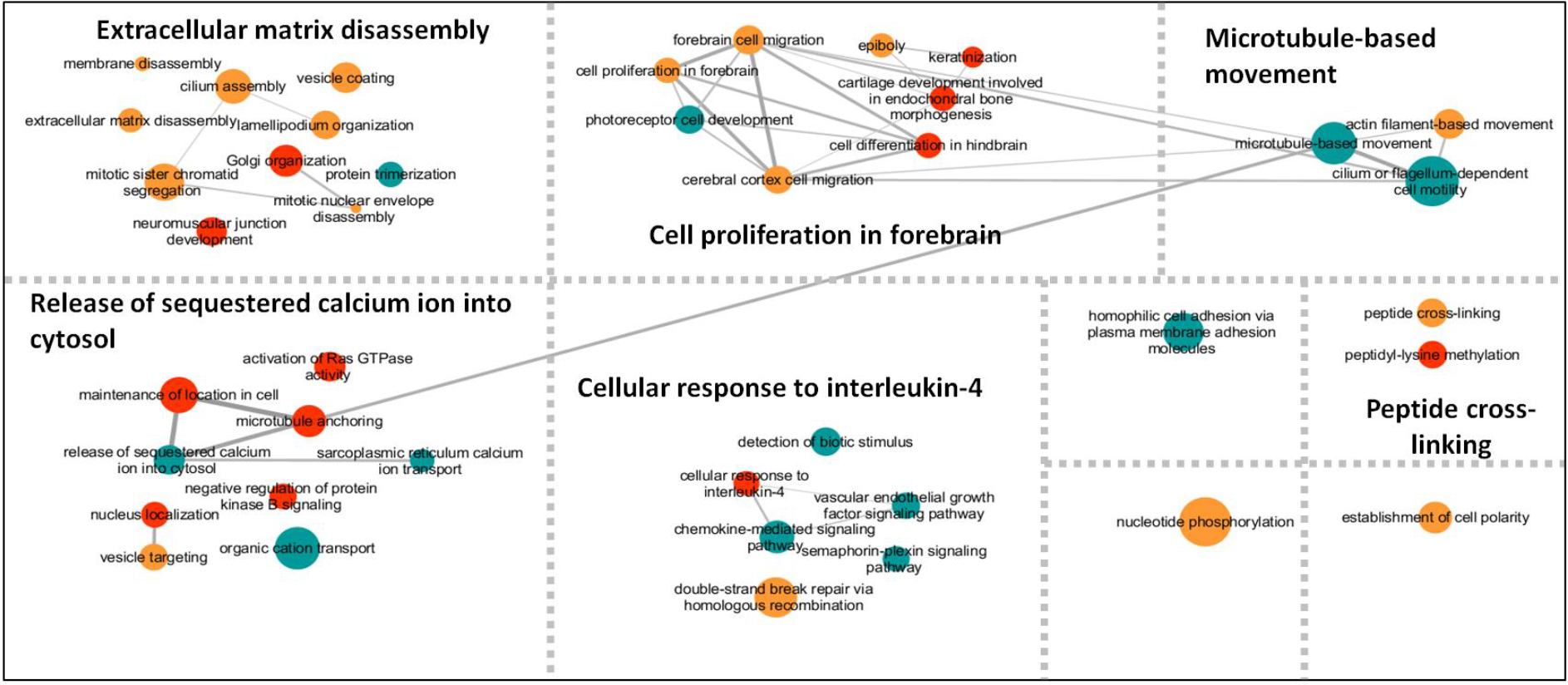
Clusters of significant GO terms enriched with variants of different frequency. Enriched GO terms were identified using three gene lists marked by variant frequency according to 1000 genomes data (novel, less than 1%, and between 1% * 5%). The resulting GO terms associated with the three gene lists are colour-coded (Cyan: between 1%%5%; Gold: less than 1%; Red: novel) and with size representing the number of genes within each GO term. The GO terms were clustered based on their functional similarity. 5 major functional categories could then be identified, namely “Extracellular Matrix Disassembly”, “Cell Proliferation in Forebrain”, “Microtubule-based Movement”, “Release of Sequestered Calcium ion into Cytosol”, and “Cellular response to interleukin-4”. Lines connecting the GO terms indicate levels of similarity between each connected pair.

## DISCUSSION

In this study, we used exome sequencing followed by Sanger validations and segregation analyses, to perform a first characterization of exome variants of likely aetiological relevance in SLI, a common form of developmental language disorder. In a dataset of 43 well-phenotyped probands, based on validation, bioinformatics characterization and previous associations, we observed potentially pathogenic variants in several genes that have already been implicated in speech-and language-related syndromes. Specifically, we identified a private start-loss variant in *ERC1*, a gene previously implicated in childhood apraxia of speech (42); a novel *de novo* substitution disrupting *GRIN2A*, a gene mutated in epilepsy-aphasia spectrum disorders (34, 57, 58); and a hemizygous disruption of *SRPX2* that was reported as causative in an earlier study of rolandic epilepsy with speech apraxia (32). Thus, although the language difficulties in SLI must (by definition) be unexpected, our findings suggest that a proportion of affected children might actually represent cases of undiagnosed developmental syndromes that may be clinically identifiable.

Consistent with accepted guidelines for defining SLI, none of the probands of our cohort were diagnosed with epilepsy. Interestingly, two of the three genes noted above were previously implicated in language-related forms of epilepsy. Disruptions of *GRIN2A* may account for between 9 and 20% of cases of Rolandic epilepsy (33, 34, 59). Coding variants affecting *SRPX2* have also been described in patients affected by Rolandic seizures, speech dyspraxia and intellectual disability, including the same variant (p.N327S) that we found in the present study (32) (but note that the relevant family was subsequently found to also carry a *GRIN2A* mutation (34)). The *SRPX2* p.N327S variant is reported to exist in control individuals with a frequency of 0.26%, although these controls were not screened for neurodevelopmental or speech and language difficulties (60). In utero silencing of rat *Srpx2* expression has been shown to disrupt neuronal migration, an effect that could not be rescued by a mutant human protein carrying the p.N327S change (61), while knockdown of the gene in mice leads to reduced vocalization (62). Clinical records did not indicate a history of seizures in our two SLI probands with variants in these genes. Our data therefore add to mounting evidence that *SRPX2* mutations may contribute to neurodevelopmental disorders in a more complex way than originally thought.

We also observed potential compound heterozygotes for putative disruptive variants of the *SCN9A* and *KMT2D* genes. *SCN9A* has been associated with febrile epileptic seizures, which themselves carry an increased risk of language impairment (63). Heterozygous loss-of-function mutations of the *KMT2D* gene are implicated in Kabuki syndrome, a severe developmental syndrome that often presents with heterogeneous oromotor, speech, and language deficits (54). The *KMT2D* variants we identified are nonsynonymous changes that may alter protein properties but are very unlikely to act as fully penetrant loss-of-function alleles, especially given that carriers of these variants do not suffer from Kabuki syndrome. Thus, if they are indeed aetiologically relevant for SLI, we must speculate that they increase risk in a subtle manner; functional assays would be required to shed more light on this hypothesis. Overall, our findings are in line with the proposed existence of shared molecular mechanisms between different neurodevelopmental disorders affecting speech and language circuits of the brain (22).

The heterogeneity of speech and language disorders and the complexity of the underlying genetic mechanisms are further illustrated by the observation that most of our cases did not carry obvious causal coding variants in known genes implicated by prior literature. We did observe novel and rare variants in candidate language-related genes in some probands, but many did not co-segregate with disorder within the family unit and their aetiological role could not be clarified. Even in cases where co-segregation was established, the small size of the family units and the limitations of phenotyping in adults limit the conclusions that can be drawn. In line with current guidelines (47), all variants would therefore require functional studies to robustly validate their relevance to SLI risk. In addition, future surveys in much larger SLI cohorts could also be informative on contributions of the various known genes to risk.

Beyond known candidate genes from the literature, we searched for variants with likely deleterious effects from elsewhere in the exome. We identified and validated two rare stop-gain variants that occurred in multiple affected children within family units. A stop-gain near the start of the *OXR1* gene was found in three siblings with speech and language-related difficulties. The OXR1 protein plays a critical role in neuronal survival during oxidative stress and is a candidate gene for amyotrophic lateral sclerosis (64). Knockout of the *Oxr1* gene in mice leads to progressive neurodegeneration and motor-coordination deficits (65). This gene therefore represents an interesting future candidate for involvement in neurodevelopmental disorder. A stop-gain in the *MUC6* gene was found in four siblings with expressive and receptive difficulties in another family. An important note of caution should be made here, since *MUC* genes are known to be particularly susceptible to false positive findings in next-generation sequencing studies, due to mapping artefacts (see http://massgenomics.org/2013/06/ngs-false-positives.html). As with all the other variants of interest that we discuss here, independent validation came from Sanger sequencing, still considered the gold standard method, which can increase confidence that these are not artefactual findings.

It has previously been postulated that some forms of neurodevelopmental disorder may follow a “double-hit” model in which combinations of events with relatively large effect sizes disrupt inter-connected pathways and substantially increase the risk of neurodevelopmental disorder (66, 67). To begin exploring this proposal with respect to SLI, we searched for genes which carried multiple rare variants of likely deleterious effect within the same proband, and probands who carried multiple events of potential interest across candidate genes. We identified several cases with multiple rare coding variants at different loci, although none occurred in genes with obvious functional connections and would thus need validation with further experimental data. One proband with multiple variants of interest carried a rare variant in the *AUTS2* gene in combination with a rare inherited haplotype in the *STARD9* gene. *AUTS2* is a long-standing candidate for autism susceptibility (68) and disruptions of this gene have been reported in individuals with developmental delay (69-72), ADHD (73), epilepsy (74) and schizophrenia (75). Indeed, it has been described as a locus that confers risk across neurodevelopmental diagnostic boundaries (43, 76). The functions of the AUTS2 protein are largely unknown but it has been suggested to play a role in cytoskeletal regulation (77). The *STARD9* gene encodes a mitotic kinesin which functions in spindle pole assembly (78). Interestingly, another proband also carried multiple rare variants in the *STARD9* gene (Figure 3). In both cases, the *STARD9* variants were not compound heterozygotes but instead appeared to represent inherited overlapping rare haplotypes that harboured multiple coding variants. The finding of co-occurring variants in two SLI probands leads us to speculate that pathways related to cytoskeletal function might be relevant for language disorders.

Potential involvement of cytoskeletal regulation in mechanisms underlying SLI susceptibility was also suggested by our independent pathway-based investigations of the exome datasets. GO analyses between and within probands converged on biological processes including microtubule-based movement, specifically the roles of dyneins and kinesins. These findings thus provide an intriguing link between the specific variants identified in single probands and the patterns of variants seen across all probands. In addition, certain biological functions appeared to cluster within variant frequency groupings. While novel and rare (0-1%) variants were over-represented within “Extracellular matrix disassembly” pathways, more common variants (1-5%) were predominantly localized within the “Microtubule-based movement” class. A potential contribution of microtubule transport pathways to risk of speech and language disorders would be of particular interest given the established links between known candidate genes for dyslexia and dynein and cilia function (79-82).

The GO categories identified as being over-represented are large functional classes. Further investigations of larger samples will be required to validate these initial findings and to elucidate whether particular subsets of genes are enriched with risk variants or whether the risk is distributed across the entire class.

The ultimate aim of exome studies is to perform an unbiased screen of all variants across the entire coding sequence. Given the sample size of the present study, we used a number of complementary methods to constrain searches for variants of interest and associated pathways. It is therefore important to note that our analyses necessarily highlight a constricted subset of loci that have supporting data from previous datasets or have an increased likelihood of aetiological significance. We have listed all identified variants within each category in the Tables presented here and as supplementary data. Nonetheless, these analyses have enabled the detection of cases with potentially pathogenic mutations (*ERC1, GRIN2A, SRPX2*), and support the role of known candidate genes and pathways (*AUTS2*, ciliary function). Moreover, our findings highlight a number of new putative candidates for future study (e.g. *OXR1, STARD9*) and novel pathways and processes (microtubule transport, cytoskeletal regulation) that may be relevant to speech and language development.

## SUBJECTS AND METHODS

### Participants

Participants for this study were taken from the SLIC (SLI consortium) cohort, the ascertainment and phenotyping of which has been described extensively in prior publications (5, 15, 46, 55, 83, 84) and were recruited from five centres across the UK; The Newcomen Centre at Guy’s Hospital, London (now called Evelina Children’s Hospital); the Cambridge Language and Speech Project (CLASP); the Child Life and Health Department at the University of Edinburgh; the Department of Child Health at the University of Aberdeen; and the Manchester Language Study. (A full list of SLIC members can be found in the Acknowledgements section.)

Briefly, the cohort comprises a set of British nuclear families who were recruited through at least one child with a formal diagnosis of SLI. This diagnosis was based on impaired expressive and/or receptive language skills (≥1.5 standard deviations (SD) below the normative mean of the general population), assessed using the Clinical Evaluation of Language Fundamentals (CELF-R) (85). The language impairments had to occur against a background of normal non-verbal cognition (not more than 1SD below that expected for their age), assessed using the Perceptual Organisation Index (a composite score derived from Picture Completion, Picture Arrangement, Block Design and Object Assembly subtests) of the Wechsler Intelligence Scale for Children (WISC) (86). Following recruitment of the proband, language and IQ measures were collected for all available siblings, regardless of language ability and DNA samples were collected from parents and children. Crucially, although there have been reports of linkage (5, 83, 84), association (15, 30, 46, 56, 87) and CNV analyses (55, 88, 89) of the SLIC families, no prior investigation has used genome-wide next-generation sequencing approaches to investigate etiology in this cohort. For the present study, we first selected unrelated probands from the SLIC cohort who had severe SLI, based on in-depth phenotypic data on multiple measures of language and cognition, along with sufficient quantities of high-quality DNA available for next-generation sequencing. This yielded a set of forty three unrelated probands for whom whole exome sequencing was carried out. The group of probands had mean scores of 65.9 (-2.3SD below expected for chronological age) and 73.8 (-1.7SD) for expressive and receptive language respectively, and a mean verbal IQ of 84.2 (-1.1SD), compared to a mean non-verbal IQ of 98.7 (-0.1SD) in line with the mean of the general population (all scores normalized to a population mean of 100 and SD of 15).

In our Figures examining family segregation of variants (see below) we present information regarding the core phenotypes; CELF-R expressive and receptive language scores, which were used to determine proband and sibling affection status. Where available, we also present data for additional phenotypes. These include the total verbal and non-verbal IQ scores from the Wechsler Intelligence Scale for Children (86) and scores on nonword repetition tasks (90). Although these were not used to ascertain affection status, they sometimes provided additional information regarding specific deficits in individuals. Nonword repetition is hypothesized to represent an index of phonological short term memory, while the IQ measures indicate general levels of verbal and non-verbal ability.

### Exome sequencing and variant discovery

Exome capture was performed using 10μg of genomic DNA from each participant. Exons and flanking intronic regions were captured with the SureSelect Human All Exon version-2 50 Mb kit (Agilent, Santa Clara, CA, USA), which is designed to capture 99% of human exons defined by NCBI Consensus CDS Database from September 2009, and 93% of RefSeq genes (~23,000). Captured fragments were sequenced using the SOLiD series 5500xl DNA sequencing platform (Life Technologies, Carlsbad, CA, USA) with 50nt, single-end runs. Sequence alignment and variant calling were performed within the GenomeAnalysis Toolkit (GATK version-2.7.2) (91). BAM files went through several stages of preprocessing, including removal of PCR duplicates using Picard Tools version-1.77 (URL:http://picard.sourceforge.net/), Base Quality Recalibration, and Indel Realignment (which form part of the GATK software package). Calling of single nucleotide variants (SNVs) was performed using a combined calling algorithm with HaplotypeCaller, which can provide a better stringency of calling and more accurate estimation of variant quality.

Raw variant calls were filtered using the Variant Quality Score Recalibration function according to GATK’s Best Practice recommendations (45), with the following training sets: human hapmap-3.3.hg19 sites, 1000G-omni-2.5.hg19 sites, and 1000G-phase1-high.confidence-SNPs.hg19 sites for SNVs, and Mills-and-1000G-gold.standard-INDELs.hg19 for INDELs. Using this training set, variant call files are recalibrated and filtered according to various parameters including the normalization of read depth (QD), the position of the variant within the read (ReadPosRankSum), the mapping quality of variant call reads (MQRankSum), strand bias (FS), and inbreeding coefficients (InbreedingCoeff). The PASS threshold after recalibration was set at 99 (99% of the testing dbSNP-137 variants could be identified using the trained model).

Filtered variants were annotated according to coordinates of human genome build hg19, RefSeq genes and dbSNP137 using the ANNOVAR annotation tool (92) which enables gene-based (e.g. functional consequence of identified changes), region-based (e.g. segmental duplications, DNAse hypersensitive sites) and filter-based (e.g. population frequencies, SIFT scores) annotations. Following annotation, all intergenic, intronic, non-coding RNAs, synonymous variants, changes that fell within a region of known segmental duplication and variants with sequencing depth below 10 in all probands were excluded from further analysis. The numbers of variants remaining at each filtering stage are shown in Supplementary Table S2. Allele frequencies were derived from 1000 Genomes Phase I (v2) data (Apr 2012) (ftp://ftp.1000genomes.ebi.ac.uk/vol1/ftp/technical/working/20120316phase1integratedreleaseversion2/) and the Exome Variants Server (evs.gs.washington.edu/esp5500_bulk_data/ESP5400.snps.vcf.tar.gz).

### Variants of potential aetiological significance: selection, validation and segregation

Since extensive investigations of exome data in individuals affected by SLI have not previously been completed, the first stage of our analyses involved the identification of sets of variants of potential aetiological significance. In accordance with current guidelines (47), we employed several complementary approaches, which considered public data sets and previously published data and employed multiple metrics followed by targeted validation and cosegregation analyses, as detailed below:

i. We considered all coding variants identified within a set of the most robust candidate genes from the literature, defined prior to the start of the analysis. This set included 19 genes (*CMIP, ATP2C2, CNTNAP2, NFXL1, FOXP1, FOXP2, DYX1C1, KIAA0319, DCDC2, ROBO1 SRPX2, GRIN2A, GRIN2B,ERC1, SETBP1, CNTNAP5, DOCK4, SEMA6D*, and *AUTS2*),as detailed in the main text.
ii. We identified rare variants (frequency of ≤0.01 in 1000 Genomes) that conferred stop-gain mutations that were predicted to be deleterious (SIFT score ≤0.05 or PolyPhen2 score ≥0.85) and that passed all the filters listed in Supplementary Table S2.
iii. We searched for potential compound heterozygotes by identifying all probands who carried two or more rare coding variants in a single gene. These variants were filtered to include only nonysynonymous or stop-gain/loss variants, splice-site changes and frame-shift INDELs that were novel or rare (frequency of ≤0.01 in 1000 Genomes (ALL) (93, 94) and the NHLBI GO ESP Exome Variant Server (EVS, ESP5400, ALL samples) http://evs.gs.washington.edu/EVS/)), and that were predicted to be deleterious (SIFT score ≤ 0.05 or PolyPhen2 score ≥ 0.85). Segregation analyses (see below) then enabled us to decipher whether the rare coding variants in the proband occurred on the same, or a different, chromosomal copy, to determine which cases were most likely to be compound heterozygotes.
iv. We highlighted potential cases of “multiple-hits” by following up all probands who had more than one variant which fell into any of the above classes of investigation. All the above variants were validated by Sanger sequencing within the probands in whom they were called. Validated variants of interest were then also sequenced in all available parents and siblings of the proband allowing the evaluation of possible segregation patterns within nuclear pedigrees.

### Pathway-based analyses

In the second stage of analyses, we performed a more exploratory investigation of biological pathways within the exome dataset. For each proband, we collated a list of all genes containing rare likely disruptive variants, defined as nonysynonymous and stop-gain/loss variants, splice-site changes and frame-shift INDELs that had a frequency of ≤1% in 1000 Genomes (ALL) (93, 94) and the NHLBI GO ESP Exome Variant Server (EVS, ESP5400, ALL samples) http://evs.gs.washington.edu/EVS/)) and that were predicted to be deleterious (SIFT score ≤ 0.05 or PolyPhen2 score ≥ 0.85). We then used the KEGG (95) and Reactome (96) databases to identify pathways affected by these variants within probands. To test whether the observed number of SLI probands sharing a particular affected pathway was higher than chance, random subject-gene associations were generated, by picking the same number of genes randomly from all genes with variants. Thus, a permuted pathway-to-subjects mapping was generated by repeating the process 1000 times. The FDR was calculated as the number of times when a pathway was seen in equal or more probands than the observed probands divided by 1000.

Following this within-proband analyses, we went on to perform gene ontology (GO) analyses in the dataset as a whole. Over-represented classes were identified across all probands using the GO database (97) and hypergeometric tests were conducted within GOstats (98) using a P-value-and FDR-level of 0.01. A list of all genes containing rare and disruptive variants (defined as above) was tested against the background gene list (all genes with all variants). FDR was obtained by multiple random sampling from the background gene list. Examinations were then split across three variant frequency gene lists; (i) genes which carried novel variants, (ii) genes that carried variants that had been reported in the 1000 genomes and EVS with a variant frequency of <1% and, (iii) as an additional comparison, we also included a list of genes which carried variants with an expected frequency of between 1% and 5%.

## ACKNOWLEDGEMENTS

DFN is an MRC Career Development Fellow at the Wellcome Trust Centre for Human Genetics, University of Oxford. The work of the Newbury lab is funded by the Medical Research Council [G1000569/1 and MR/J003719/1]. Core services at the Wellcome Trust Centre for Human Genetics are funded by Wellcome Trust (grant reference 090532/Z/09/Z) and MRC (Hub grant G0900747 91070). This study was supported by the Max Planck Society. We are grateful to Christian Gilissen for helpful comments on the manuscript.

Members of the SLI Consortium: Wellcome Trust Centre for Human Genetics, Oxford: D. F. Newbury, N. H. Simpson, F. Ceroni, A. P. Monaco; Max Planck Institute for Psycholinguistics, Nijmegen: S. E. Fisher, C. Francks; Newcomen Centre, Evelina Children’s Hospital, St Thomas’ Hospital, London: G. Baird, V. Slonims; Child and Adolescent Psychiatry Department and Medical Research Council Centre for Social, Developmental, and Genetic Psychiatry, Institute of Psychiatry, London: P. F. Bolton; Medical Research Council Centre for Social, Developmental, and Genetic Psychiatry Institute of Psychiatry, London: E. Simonoff; Salvesen Mindroom Centre, Child Life & Health, School of Clinical Sciences, University of Edinburgh: A. O‘Hare; Cell Biology & Genetics Research Centre, St. George's University of London: J. Nasir; Queen’s Medical Research Institute, University of Edinburgh: J. Seckl; Department of Speech and Language Therapy, Royal Hospital for Sick Children, Edinburgh: H. Cowie; Speech and Hearing Sciences, Queen Margaret University: A. Clark, J. Watson; Department of Educational and Professional Studies, University of Strathclyde: W. Cohen; Department of Child Health, the University of Aberdeen: A. Everitt, E. R. Hennessy, Shaw, P. J. Helms; Audiology and Deafness, School of Psychological Sciences, University of Manchester: Z. Simkin, G. Conti-Ramsden; Department of Experimental Psychology, University of Oxford: D. V. M. Bishop; Biostatistics Department, Institute of Psychiatry, London: A. Pickles

